# Savanna-forest dynamics: Encroachment speed, model inference and spatial simulations

**DOI:** 10.1101/2024.03.12.584640

**Authors:** Yuval R. Zelnik, Ivric Valaire Yatat-Djeumen, Pierre Couteron

## Abstract

Forest encroachment over savannas has been recurrently reported in the tropics over the last decades, especially in northern tropical Africa. However, process-based, spatially-explicit modelling of the phenomenon is still trailing broad scale empirical observations. In this paper, we used remotely-sensed diachronic data from Central Cameroon to calibrate a simple reaction-diffusion model, embodying dynamical interactions between grass and woody biomasses in the savanna biome. Landsat satellite image series over the Mpem and Djim National Park witnessed a dramatic extension of forest over the last five decades and our estimates of forest front speeds based on randomly sampled transects indeed yielded higher values (5-7 meters per year) than in the existing literature. We used simulations of the model to provide the first hitherto estimates of woody biomass dispersal coefficients. Since the region under study did not provide examples of savanna progression, estimates of grass dispersal proved inconsistent and we reverted to literature-based historical data to reach rough estimates. This paper demonstrates that broad scale remote sensing data allows for calibrating simple reaction-diffusion models of vegetation dynamics in the savanna biome. Once calibrated, such models become a general baseline of expected changes and a valuable tool to understand how spatial environmental factors (e.g., soil substrate) may locally modulate the overall dynamics.

## 2 Introduction

Savannas and grasslands cover a substantial portion of the global land surface, particularly in sub-Saharan Africa, and play a crucial role in supporting local populations and biodiversity (Parr *et al*., 2014; Aleman & Staver, 2018). Grasslands are dominated by grasses with no or very weak woody cover, while savannas are characterized by a mixture of woody species, grasses, and forbs, with a discontinuous tree canopy and a continuous grass understory.

The maintenance of savannas relies on specific mechanisms and disturbances that contribute to their open canopy structure and species diversity (Scholes & Archer, 1997). As soon as climate is sufficiently moist (e.g. above 1000 [mm/yr], (Staver *et al*., 2011)), savannas require frequent disturbances, primarily through fires and/or mega-herbivores, which restrain the encroachment of trees and help maintain an ecosystem with a mixture of trees and grasses. Fires reduce tree density, promote grass growth, and prevent the establishment of a closed forest canopy (Beckett *et al*., 2022). Such disturbances create a favorable environment for grasses to thrive and ensure the coexistence of tree and grass species, the later being more adapted to high-light environments, (Smit & Prins, 2015).

The occurrence and stability of savannas thus depends not only on climate, but is also influenced by feedback mechanisms between fire, canopy cover, vegetation, soils and hydrology, in addition to grazing and direct human intervention in these processes (Scholes & Archer, 1997; Smit & Prins, 2015). In Africa, while forest overwhelmingly dominates for rainfall over 1500 [mm/yr], both forests and savannas can persist in intermediate rainfall regimes (1200-1500 [mm/yr]) with a short dry season (under 4 months), sometimes in the same landscapes, suggesting they may represent alternative stable states. Positive feedback between fire and canopy openings (’grass-fire feedback’) plays a role in maintaining these states. Savannas are characterized by a unique and highly biodiverse functioning, offering multiple ecosystem services. However, ongoing land use changes, such as large-scale conversion to commodity crops and forest plantations, pose threats to savanna ecosystems, emphasizing the importance of implementing appropriate management strategies that consider the specific needs of savannas and their ecological dynamics.

In addition, forest encroachment into grasslands and savannas is a global trend that is significantly altering the landscape structure and ecological dynamics of these ecosystems (Stevens *et al*., 2017). Forests, savannas and grasslands often form a complex matrix in tropical climates, creating diverse habitats that support local livelihoods and foster biodiversity (Parr *et al*., 2014). However, recent observations indicate a rapid shift, with forests increasingly encroaching upon savannas in regions such as sub-Saharan Africa, homogenizing the formerly-complex forest-savanna mosaic (Mitchard *et al*., 2011; Venter *et al*., 2018; Stevens *et al*., 2017)

Forest encroachment is a multifaceted phenomenon that is driven by various factors, including climate change, human activities, and changes in species composition (Stevens *et al*., 2017; Venter *et al*., 2018; Devine *et al*., 2017). Despite its growing importance, there is a lack of understanding of how much and how fast forest encroachment is happening, what factors influence it, and what the underlying mechanisms are (Devine *et al*., 2017). Addressing these questions is crucial for improving our understanding of forest-savanna dynamics and for informing conservation and land-use planning.

The technical and conceptual challenges associated with studying forest encroachment are daunting. They include very large spatial and temporal scales of the ecosystem and the encroachment process overall, difficult logistical and political situations in many regions, and above all a lack of a unifying perspective. Moreover, the scale and pace of forest encroachment make it difficult to observe and quantify the effects of specific drivers on the landscape.

Several tools have been used to understand savanna dynamics and the encroachment of forests into them. Local scale field data (measuring trees, soil properties) have been used to explore the dominant processes in these ecosystems (Guillet *et al*., 2001; Mitchard *et al*., 2011), but they are difficult and expensive to conduct over long time spans, and are rarely done on large scales that contain entire landscapes. Remote sensing at large spatial scales (satellite images, aerial images) are particularly useful tool to capture encroachment (Stevens *et al*., 2017; Venter *et al*., 2018; Mitchard *et al*., 2011; Sagang Takougoum *et al*., 2022), with the main drawback being the limited availability of data earlier than the past five decades.

Dynamical models can overcome some of the drawbacks of limited data, as they can allow us to extrapolate beyond the existing data, bridge the scale gap between field and remote-sensing data as well as attempt to predict future behavior. In the context of forest and savanna interactions, most models are spatially implicit, assuming spatial homogeneity, and therefore they cannot capture spatial processes (Staver & Levin, 2012; D’Onofrio *et al*., 2015; Yatat *et al*., 2021). Spatially explicit models are less often used, and mostly fall into two categories. The first are Cellular Automata models (Favier *et al*., 2004a), which although conceptually simple are difficult to parameterize and analyze. More recently, spatially explicit models using partial differential equations (PDEs) have been used (Tchuinte *et al*., 2014; Yatat *et al*., 2018; Tega II *et al*., 2022), which are more easily parameterized and can be analyzed using mathematical tools.

Here we use a combination of remote-sensing results (Sagang Takougoum *et al*., 2022) and dynamical models to assess forest encroachment, considering a case study in the forest-savanna transition zone of Cameroon, central Africa. We also demonstrate how, using a spatially explicit modeling approach where we simply add diffusion terms to the minimalistic temporal model of Yatat *et al*. (2021), can give valuable and novel insight on forest-savanna dynamics.

## 3 Methods

### 3.1 Model

To model the interactions between forest and savanna, and in particular forest encroachment, we use an extension of the model by Yatat *et al*. (2021). The original model in its simpler form follows the dynamics of trees *T* and grasses *G* above-ground biomasses in a non-spatial form, using two coupled ordinary differential equations (ODEs). We extend this model, similarly to Yatat *et al*. (2018), by adding spatial diffusion terms for both variables, resulting in two coupled partial differential equations (PDEs). This new model is spatially explicit, allowing us to explore forest encroachment by front propagation, and its interaction with spatial heterogeneity of the landscape. In our model, we consider the savanna landscape to include grasslands (i.e. regions with only grasses) as well, and we henceforth use the term savanna alone to describe any grass-dominated habitat. The model reads:

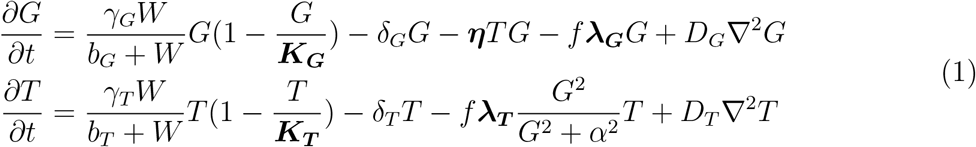

With the following definitions of the terms in bold:

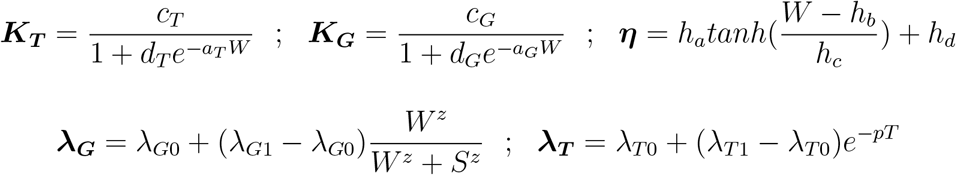

The parameters and estimates of value ranges (from Yatat *et al*. (2021)) are given in table 1.

**Table 1:**
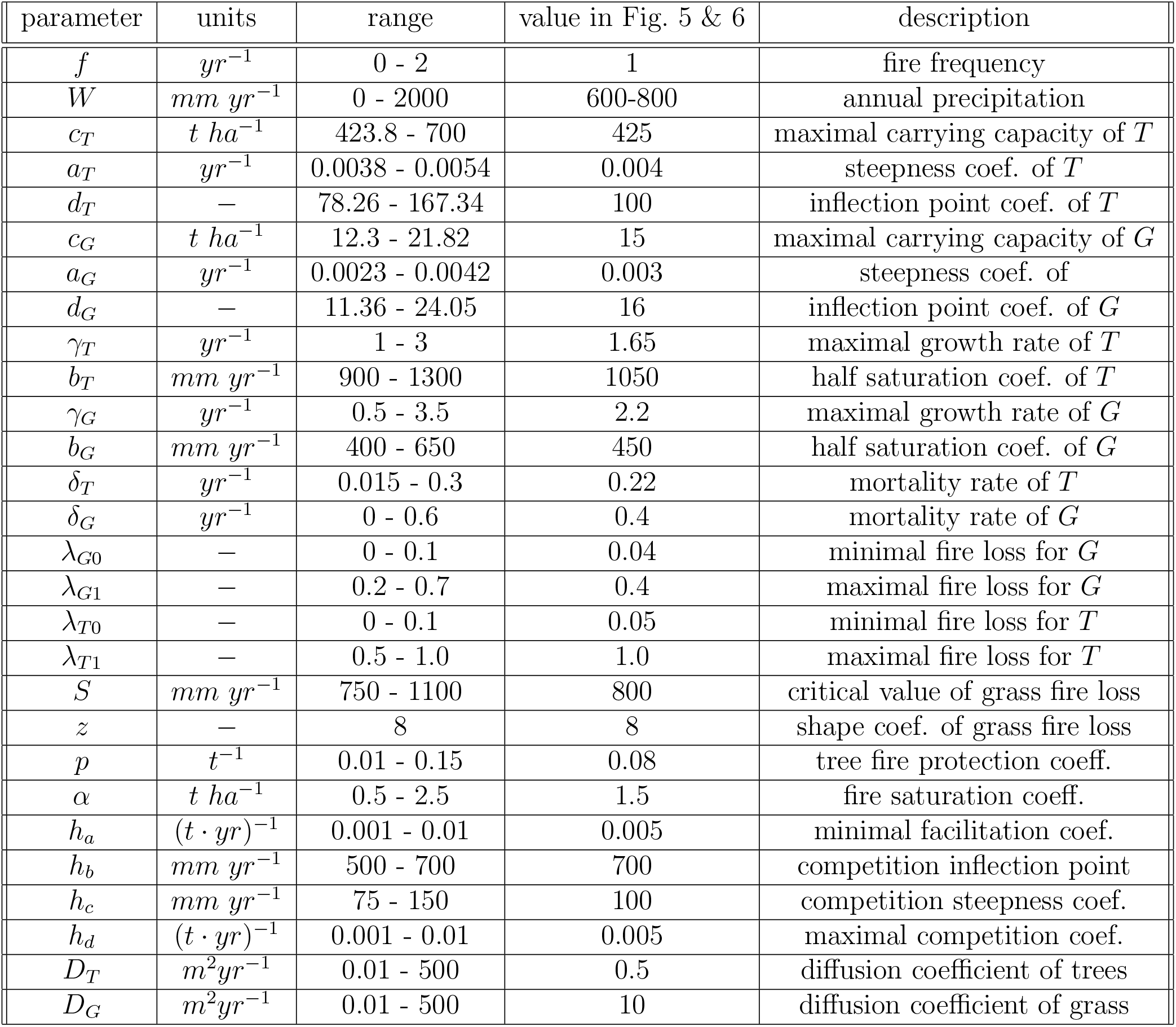
The model includes 28 parameters in total. We introduce two spatial parameters, *D*_*G*_ and *D*_*T*_. For the other 26 parameters, their ranges are taken from Yatat *et al*. (2021).

In its non-spatial form (Yatat *et al*., 2021), this model has four steady-state solutions: bare-soil (*T* = *G* = 0), forest (*T >* 0; *G* = 0), savanna (*T* = 0; *G >* 0), and mixed (*T >* 0; *G >* 0). Depending on parameters, different combinations of these solutions may be linearly stable, including a bistability between forest and savanna, and a tristability of the forest, savanna and mixed states (Yatat *et al*., 2021).

### 3.2 Parameter Inference

We estimate the parameter values for the two parameters related to spatial processes – *D*_*T*_ and *D*_*G*_. To do this, we randomly choose values for the other 26 parameters within their given ranges, and infer on the most likely value of the two diffusion coefficients, given an observed front speed.

The inference is done by using a set of *n*_*p*_ different viable parameter sets (each set has a given value for the 26 parameters, i.e., excluding *D*_*T*_ and *D*_*G*_), and *n*_*o*_ different observations of front speeds. This gives a total of *n*_*p*_*n*_*o*_ of points from which to infer on the likely distribution of diffusion coefficients.

The *n*_*o*_ observations of front speeds are a result of an estimation of front speeds from remote sensing data (see more details in the next subsection). The *n*_*p*_ parameter sets are a result of the following two steps: i) 1000 parameter sets are chosen at random, where each parameter value is taken from a truncated Gaussian distribution, with 95% of the distribution falling inside each parameter’s value range (see table 1), and a minimum value is taken as 10% below the low value of the range (e.g., the minimum value for *γ*_*T*_ is 0.1). ii) We choose a subset of these 1000 parameter sets, according to the equilibrium solutions. We test two options, one where we require that both the forest and savana solutions exist, and at least one of them is stable, and the second where we require that both solutions are stable. This gives us *n*_*p*_ of 869 and 186, respectively. In the main text we focus on the former option, i.e. not requiring bistability, since the results of the two options are similar (See Appendix).

For a specific parameter set and estimation of front speed, the inference of diffusion parameters is done in the following way. A wide range of different values for the diffusion parameters are tested, with 48 values between 0.01 and 500, equally spaced on a logarithmic scale, giving 2304 pairs of values (of *D*_*G*_ and *D*_*T*_). For each pair, a simulation of a front between a forest and a savanna state is simulated with the specific parameter set, in a system of one spatial dimension. From this simulation the front speed is estimated numerically, giving 2304 values of front speeds. The best pair is then chosen, as having a front speed that is closest to the observed front speed, yielding an estimation of *D*_*T*_ and *D*_*G*_. If one or more simulated front speeds is within a 1% error from the observed speed, we choose one low-error pair at random (to avoid artifacts). We do not keep any pair if no simulation could reproduce a similar front speed (i.e. if all simulations have above 1% error). Repeating this *n*_*p*_*n*_*o*_ times results in a distribution of inferred diffusion parameters.

### 3.3 Front Speed Estimation

To estimate the front speed of forest encroachment, we focus on a case study of the forest-savanna mosaic in Mpem and Djim national park in Cameroon (MDNP, (Sagang Takougoum *et al*., 2022); see Fig. 1). The MDNP harbours mosaic landscapes entailing close canopy forests and very open savannas of low woody biomass and very high biomass of heliophytic Poaceae. Average total annual rainfall is of ca. 1350 mm with a short dry season extending from December to March (Sagang Takougoum *et al*., 2022). Using remote sensing data from the Landsat satellite series over the span of 43 years (from 1978 to 2020), we designate each pixel of 30×30 [m] into one of 11 categories (with integer values from 0 to 10). The two extreme categories of savanna or forest cover, (observable in 2020) and nine intermediate categories of savanna turning into forest at different points in time. The national park experienced a swift and regular increase in forest cover over the monitored period, from ca. 30 to 70 percent of its total area (see Sagang Takougoum *et al*. (2022) for details). Conversely, forest boundary recession was not observable. With this spatio-temporal data, we can find multiple instances of forest encroachment into savanna, using an algorithm that randomly finds relevant locations for these dynamics. This algorithm is detailed below, and its end result is a set of *n*_*o*_ transects along which we can observe the forest encroaching into the savanna. From each transect we extract a profile of a moving front, with which we can estimate the encroachment speed, thus giving us *n*_*o*_ front speed estimates.

**Figure 1:**
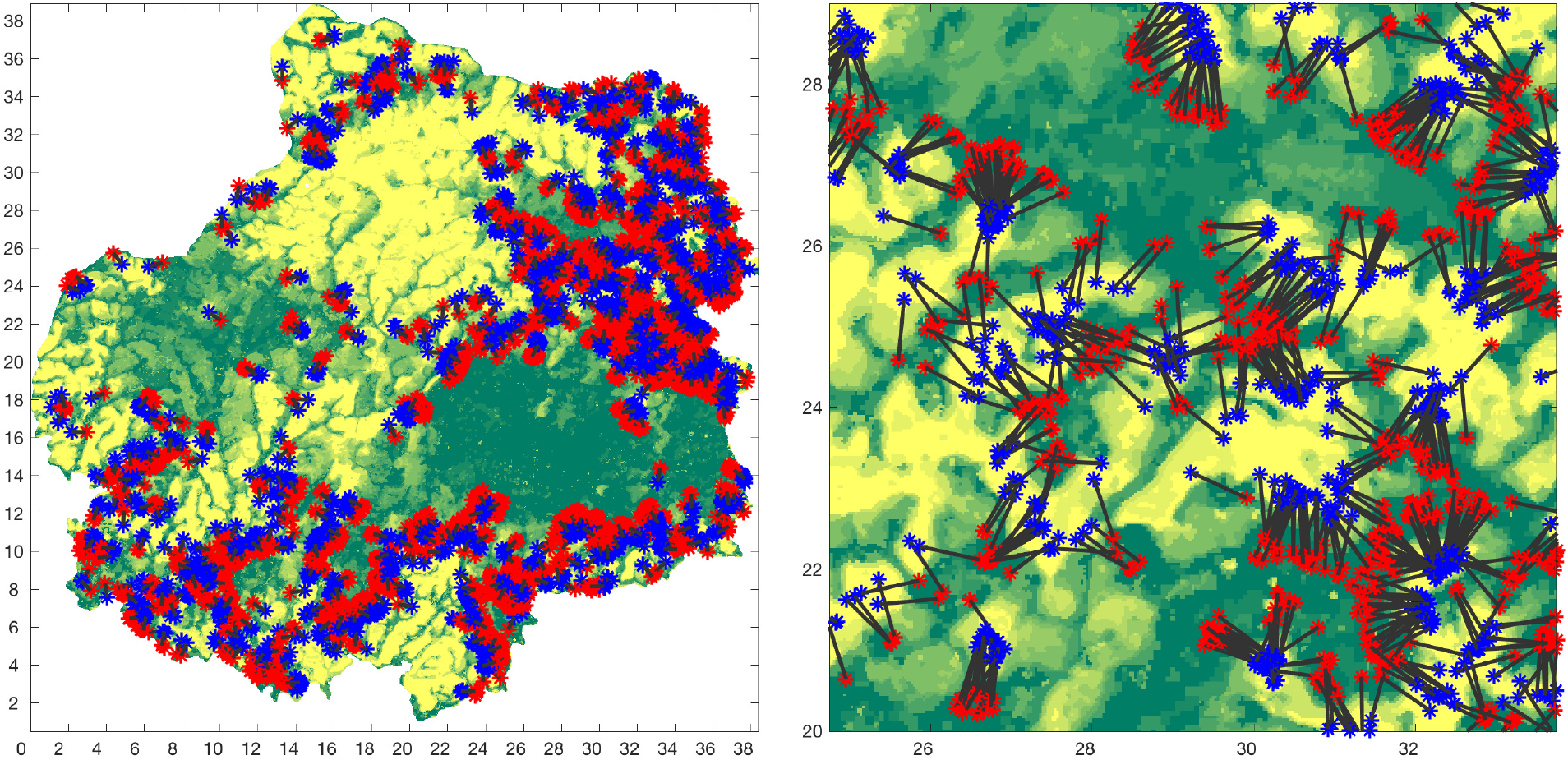
Locations of assessment of front dynamics, overlayed on the map of the Mpem and Djim national park (MDNP) in Cameroon. Left panel shows the entire park region, while right panel shows a small region in more detail. Shades correspond to different states of the landscape, with dark green corresponding to forest cover in the past 50 years, light yellow to locations with savanna cover to this day, and shades in between to locations where forest has taken over savanna at different times. Locations where a forest-savanna front structure was found (see details in methods) are shown by red and blue asterisks (corresponding to the forest and savanna edges of the front, respectively) connected by a black line.

#### 3.3.1 Finding front transects

The algorithm for finding transects along which we observe forest encroachment is made up of the following four steps (Fig. 2, left panel): 1) randomly choose a focal pixel within the temporal image. If it is a forest pixel proceed to the next step. 2) Test if enough pixels in its immediate vicinity are also forest. If more than a fraction of *c*_*f*_ pixels within the radius *c*_*r*_ from the focal pixel are forest, proceed to the next step. 3) Choose a pixel at a distance of *c*_*d*_ from the focal pixel, with a random angle. If the pixel is savanna and in its vicinity of *c*_*r*_ there are at least a fraction of *c*_*f*_ savanna pixels, proceed to the next step. Otherwise, repeat this step up to *c*_*n*_ times (each time choosing a different angle at random), until either a good savanna pixel with savanna surroundings was found, or, if *c*_*n*_ attempts were made, we discard this attempt and go back to step 1. 4) Save the locations of the focal pixel (forest) and the secondary pixel (savanna), to be used later to extract a profile of a moving front. These steps are repeated 10^6^ times. For the algorithm, we use *c*_*f*_ = 0.85, *c*_*r*_ = 4 to makes sure that the focal and secondary pixels each represent a cluster of forest and savanna, respectively. We use *c*_*n*_ = 20 to make sure we reasonably tested all the region surrounding a focal pixel. We use 7 different values of *c*_*d*_, from 10 to 40 in regular spacing, and for each one we run the the aforementioned algorithm 10^6^ times, and coalesce all the transects found.

**Figure 2:**
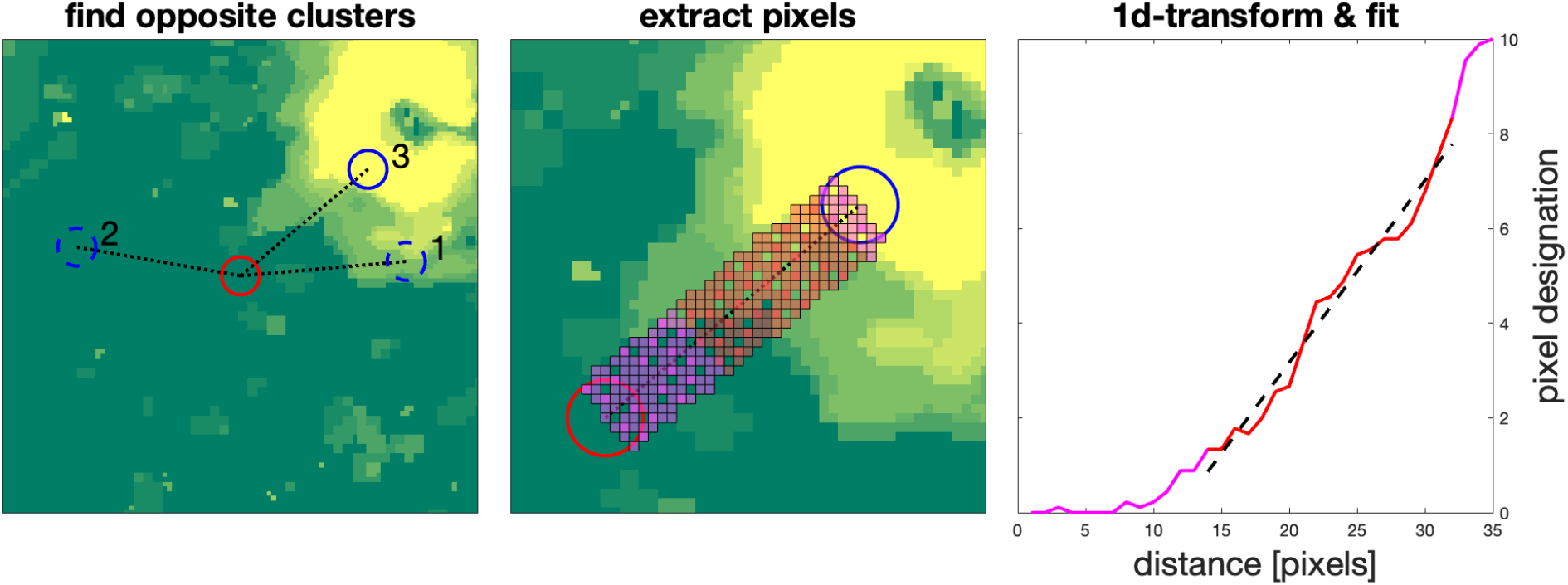
Demonstration of finding a transect and estimating its front speed. Left panel shows an initial forest cluster found (red circle), and three attempts at finding an opposite savanna cluster at random angles (blue circles). The first two clusters found are discarded (dashed circle) as they do not contain savanna, while we continue with third one found (solid circle) as it does contain savanna. Middle panel shows how a transect region is extracted between the two clusters (shown in blue and red circles, same as left panel). Pixels used are shown by overlaid red and magenta color. Each line of pixels that is perpendicular to the dotted line is averaged to one value, shown in the right panel. The result, shown in the right panel, is a curve going between 0 (forest) to 10 (savanna), showing the dynamics of a front. The region used for a linear fit is shown in red, while the rest is shown in magenta (colors correspond to the colored pixels in middle panel). From the linear fitted function (dashed black line), we use the slope to calculate the front speed. Each pixel in the left and middle panels is 30[m]x30[m].

#### 3.3.2 Estimating front speed from transects

Given two locations, one of a forest cluster and the other of a savanna cluster, we can estimate the speed of a front along the transect that connects these two points. We do this by first extracting a one-dimensional transect from the image (Fig. 2, middle panel), and then estimating the front speed along this transect by fitting a linear function (Fig. 2, right panel).

To extract the transect, we define the transect length *c*_*l*_ as the distance in pixels between the two clusters (one of forest, one of savanna), and the transect width *c*_*w*_ = 2*c*_*r*_ = 8 as the number of pixels in the perpendicular direction to the line connecting the two clusters, that defines the transect. Together this makes a rectangle of dimensions *c*_*l*_ and *c*_*w*_ (in pixels) that connects between the two clusters (Fig. 2, middle panel). To extract the transect, we collapse the rectangle into a line of length *c*_*l*_, so that the end result is a vector of length *c*_*l*_, with each point in this vector being an average of *c*_*w*_ pixels within the rectangle, taken along the perpendicular line to the line connecting the two clusters.

With the result of the transect extraction, a transect profile of *c*_*l*_ points, we can now estimate the front speed – the speed at which the forest encroachment occurs. To do this, we first throw out the points with extreme values on both sides of the profile, in our case values below 1 and above 9 (where the extreme most values are 0 and 10). This is to prevent artifacts and focus on the forest encroachment itself. We then fit the remaining points in the profile to a linear function. Using the slope of this fit *ϕ* (where a slope of 0.5 means that 2 pixels are traversed every 1 time unit, translating to 2×30=60 meters within 5 years), we now estimate the front encroachment speed as: *v* = 30*/*(5*ϕ*) = 6*/ϕ*[*m/yr*]

#### 3.3.3 Evaluating quality of transects

We score transects using five different quality measures, and retain the front speed estimates only of those transects that have good scores in all five measures. The measures are: *m*_1_: homogeneity of forest cluster; *m*_2_: homogeneity of savanna cluster; *m*_3_: one-dimensionality of transect; *m*_4_: monotonicity of profile; *m*_5_: regularity of profile. The first two measures test the homogeneity of the two clusters (of forest and savanna, respectively). The third measure tests if the transect is taken roughly perpendicular to a simple front between two domains. The fourth and fifth measures look at the mid-product of the profile, and test if it is monotonous, i.e. if it shows a steadily increasing trend of forest encroachment, and how regular is this trend, respectively. The goal of these measures is to ensure that our front speed estimation can be compared to a one-dimensional system and that landscape heterogeneity plays a minimal role. That is, we target regions where there is a clear line separating between forest and savanna, and where the propagation of the forest into the savanna occurs perpendicular to this line.

*m*_1_ and *m*_2_ are enacted by our choice of clusters (see above), and defined as the fraction of non-forest and non-savanna pixels in the radius *c*_*r*_ in the first and second clusters, respectively. *m*_3_ is calculated by estimating the standard deviation of normalized values (i.e. values divided by the overall range) in the perpendicular direction to the transect, averaged along the transect. *m*_4_ is the largest change in the opposite direction to that of the front, normalized by the overall range. *m*_5_ is the root-mean-square error of the linear fit of the profile. We keep only clusters for which *m*_*i*_ *< c*_*x*_ for *i* = 1, 2, 3, 4, 5, with *c*_*x*_ the threshold value of 0.6, 0.4, or 0.2.

## 4 Results

We show here the estimates of front speed using our case study, followed by inference of diffusion coefficient values for a spatially-explicit model. Finally we demonstrate how a spatially-explicit model can provide insight on the dynamics of forest encroachment and its interaction with the landscape template.

### 4.1 Estimation of front speed

Using the case study of MDNP (Fig. 1), we arrive at estimations of the front speed at which forest encroachment is taking place (Fig. 3). We show the estimations for different conditions, where we retain different number of transects according to their quality (see methods). Overall the results are not highly sensitive to the threshold chosen, with a tendency for slower speed estimation if we retain only the best transects, i.e. transects with the best scores. We note that estimates of front speed below approximately 1 [m/yr] are not possible given the spatio-temporal resolution of our data. Overall, our results suggest a median front speed of about 6 [m/yr], but with a range of values mostly between 2 and 12 [m/yr].

**Figure 3:**
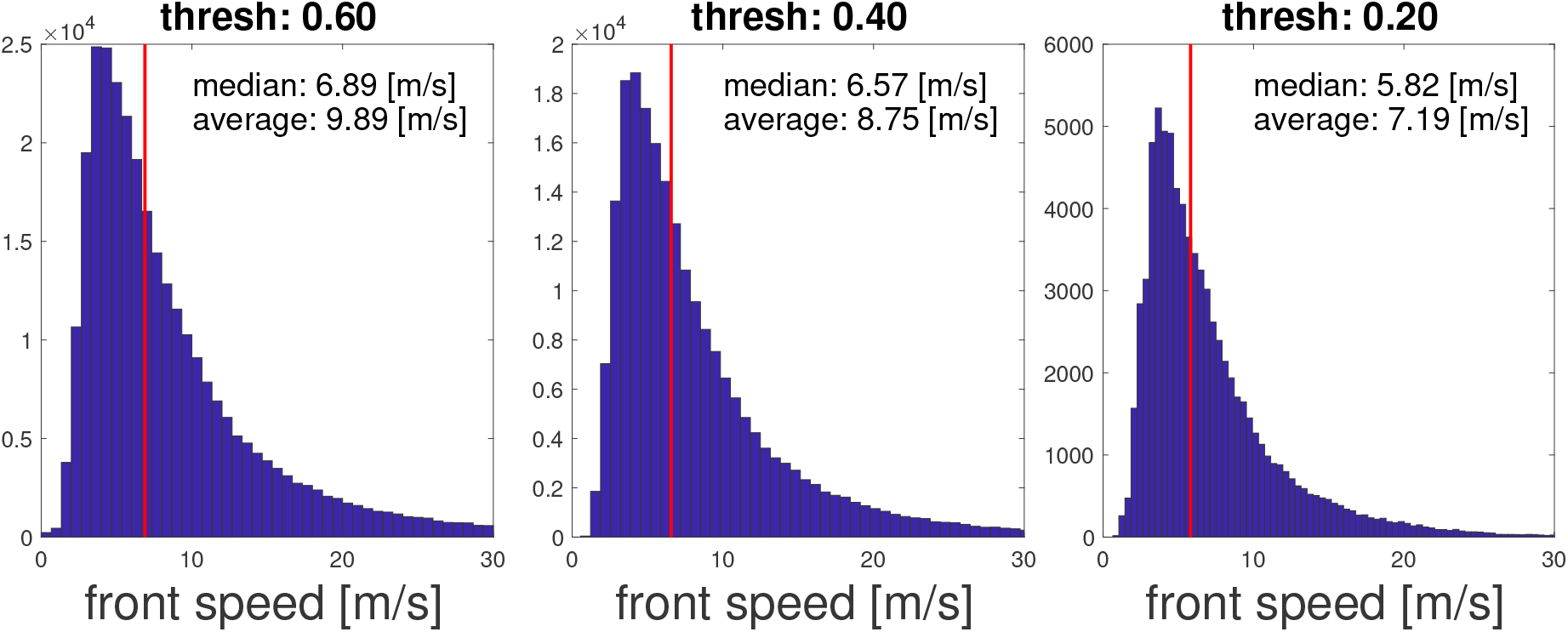
Estimates of forest encroachment front speed. The histograms show estimated values of front speed, derived from remote sensing images of MDNP. Median and average values are noted, and median value is also shown with a vertical red line. Different panels correspond to speed estimates that score below a threshold of 0.6, 0.4, and 0.2, for all five measures considered (see details in methods). This results in 306992, 217231, and 78935 estimates, respectively. For clarity, estimates of speeds above 30 [m/yr] are not shown (but are used for calculations).

### 4.2 Inference of model parameters

To infer the diffusion parameters *D*_*T*_ and *D*_*G*_, we compare a distribution of front speeds based on field data with front speeds from simulations (see Methods). Using the case study of MDNP (Fig. 1) for estimation of front speeds (Fig. 3), we can compare these speeds with model results. For tree spread (*D*_*T*_), we reach a distribution shown in the left panel of Fig. 4, with median value of 10 [m^2^/yr]. 90% of values lie between 1 and 100 [m^2^/yr]. Estimating grass spread (*D*_*G*_) in a similar fashion does not work as well, since model results of front speed are generally not sensitive to *D*_*G*_ when the forest is encroaching, as observed in MDNP. Given that all our data shows forest encroachment, inference of *D*_*G*_ is not possible with this method.

**Figure 4:**
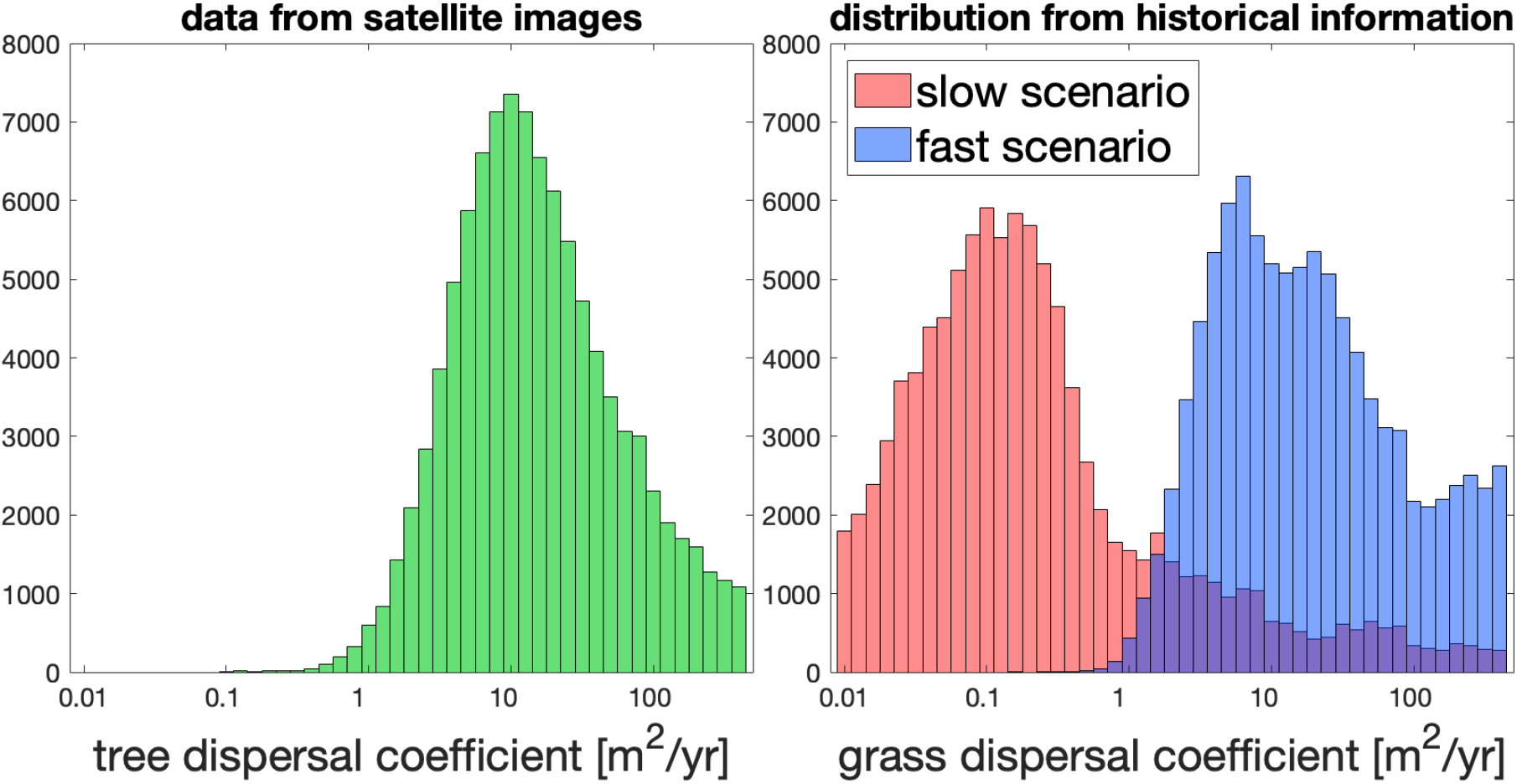
Inference of model parameters: diffusion coefficients *D*_*T*_ and *D*_*G*_. Left panel shows *D*_*T*_ inference when confronting model results with front speed estimates from satellite images. Right panel shows *D*_*G*_ inference based on rough estimates of possible historical scenarios of grass expansion in the late Holocene.

As a rough substitute, we look at historical data on the region, as analyzed by Lebamba *et al*. (2016). Based on these estimates and extrapolation of savanna and forest spatial extent in MDNP, we consider two possible scenarios of historical savanna spread, which we term fast and slow (see more details in Appendix). For these two scenarios we build log-normal distributions for front speeds, with median values of 0.25 [m/yr] and 2.65 [m/yr], for the slow and fast scenarios, respectively. With these slow and fast scenarios we reach the inference shown in the right panel of Fig. 4, with median values of 0.16 and 15.8 [m^2^/yr], respectively. These values can be understood as defining the likely range of *D*_*G*_ values we may expect to see, and indeed they are in line with other estimates in the literature (Zelnik *et al*., 2015).

### 4.3 Exploring mechanisms of forest encroachment

The strength of process-based models with diffusion terms is that they allow for predictions on boundary movements and landscape dynamics while explicitly taking into account pre-existing spatial structures in environmental heterogeneity (e.g. in soil or topography). The determination of front speed (Fig. 3) and the use of spatially explicit models (for which we estimated the parameters in Fig. 4) is useful when considering the spatial nature of forest encroachment. We illustrate here two ways in which we may need to explicitly consider spatial structure, and in particular use a two-dimensional spatial representation.

The most straight-forward way in which spatial structure can impact the effective rate of spatial processes such as forest encroachment, is by creating obstacles for spread. This is demonstrated in Fig. 5 (bottom), in which we assume some part of the region shown is more constraining to growth due to lower water availability (e.g. due to different soil). Comparing the top and bottom panels, we can see that spread of the forest (green) is significantly faster when there are no such obstacles (top panels). We note that in a one-dimensional setting this effect cannot be represented since such an obstacle cannot be overcome.

**Figure 5:**
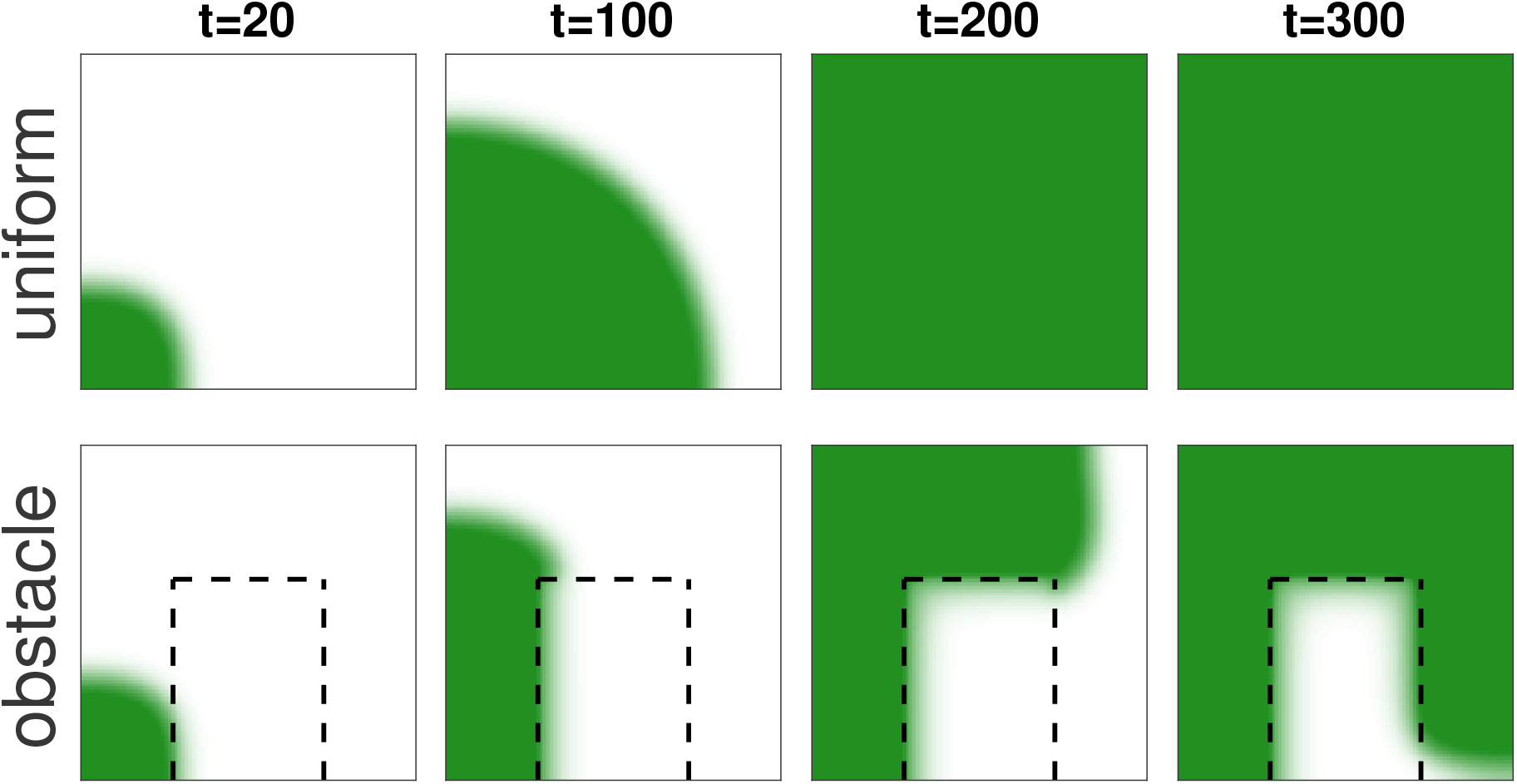
Impact of spatial structure on overall encroachment speed. Darker green color show higher density of trees (white equals no trees). On the top panels a region in the center is drier (lower values of *W*), leading to an overall slower forest encroachment. While on the top panels the encroachment is complete by *t* = 200, on the bottom panels it is still ongoing at *t* = 300. All parameters as defined in Table 1. We set *W* = 800, except for the rectangle of lower water availability (*W* = 600) in bottom panels. Time is in units of years.

Additionally, the shape of spatial heterogeneity can also affect front movement. In Fig. 6 we compare how forest encroachment is affected by two different shapes of spatial heterogeneity, due to curvature effects. On the top panel, a straight shape of the drier region leads to a slow-down of the forest encroachment. However, the bottom panel shows that a circular shape leads to a further slow-down of encroachment, so that effectively the encroachment has stopped.

**Figure 6:**
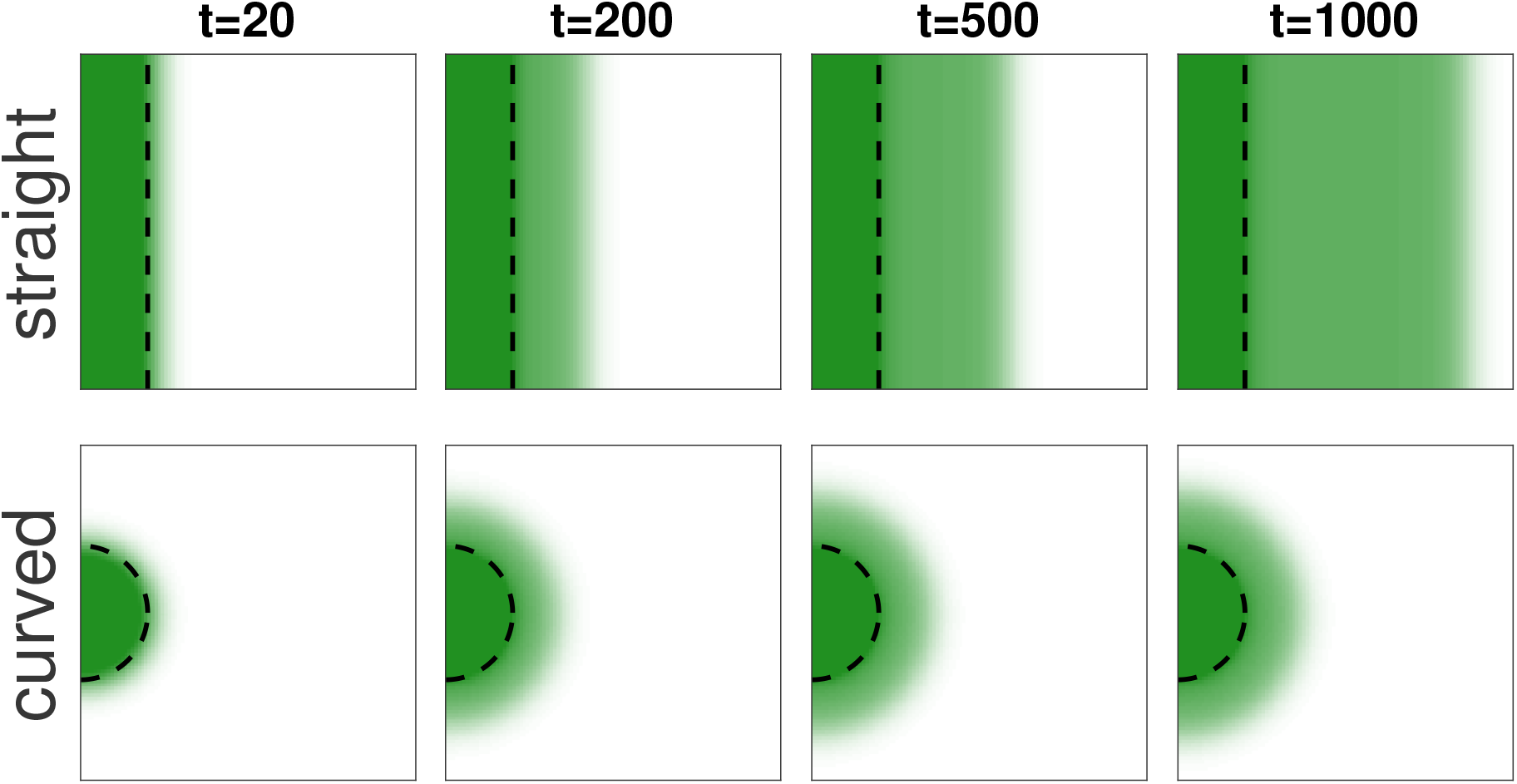
The effect of curvature on front speed. Darker green color show higher density of trees (white equals no trees). Both top and bottom panels contain drier regions, which the front reaches around *t* = 20. While an encroachment is seen at the top (compare *t* = 1600 to *t* = 200), virtually no encroachment is being seen at the bottom panels. All parameters as defined in Table 1. We set *W* = 800 except for the drier regions (*W* = 700), to the right of the black dashed line in each panel. Time is in units of years.

## 5 Discussion

We conducted a detailed estimation of forest encroachment front speed (Fig. 3), revealing higher rates than previously observed in similar ecosystems (Schwartz *et al*., 1996; Favier *et al*., 2004b). Using these speed estimates we calculated diffusion parameters within our model (Fig. 4). We further show that employing such parameterized dynamical models is a valuable tool, due to the inherent complexity of landscapes. We demonstrate the specific relevance of dynamical spatial modeling in two-dimensional space, by highlighting phenomena such as obstacle avoidance (Fig. 5) and curvature effects (Fig. 6).

We found substantial variability in both front speed estimates as well as in diffusion parameter estimates. Given that tighter thresholds (i.e. cleaner fronts) yield narrower speed distributions (moving from left to right in Fig. 3), it appears likely that much of this variability is related to the estimation method and fine-scale heterogeneities in the landscape. This in turn suggests that in homogenous conditions the spread rate is probably relatively well defined. Moreover, median speed estimates do not appear to vary much (compare panels of Fig. 3, and as such are useful for characterizing the landscape dynamics. The variability in estimation of model parameters is further exasperated by our uncertainty of other parameter values. In its current state, it is mainly useful for rough estimation of model parameters, for which we have little if any available estimates in the literature. Refining these estimates will likely require either better estimates of other model parameters (e.g. using small scale experiments) or finding other spatial patterns which we can use as confounding factors for our estimations.

It is insightful to compare our encroachment speed estimate to other similar estimates in the literature. In Central Africa, comparable situations reported slower encroachments speeds, of 0.5-2 [m/yr] (Favier *et al*., 2004a; Youtta, 1998), compared to our estimates of 5-7 [m/yr] (Fig. 3). It is possible that past estimates of slower speeds were constrained by the the method employed – often these were based on a limited number of transects. Larger scales estimates typically focus on overall cover change, e.g. as a fraction of the region, and are thus not as comparable (Mitchard & Flintrop, 2013; Stevens *et al*., 2017).

Spread speeds can vary widely across different ecological contexts. Rapid spread rates of kilometers per year have been observed in single-species invasions and range shifts (Caswell *et al*., 2003; Phillips *et al*., 2006; Chamorro *et al*., 2020). In contrast, the slower spread rates are observed in ecosystems experiencing spread movements, i.e. the movement of the ecotones. Front speeds observed include less than 1 [m/yr] for woody encroachment in drylands (Drees *et al*., 2023) or 1-100 [m/yr] in boreal range expansion (Rees *et al*., 2020), likely reflect the interplay between species dynamics and environmental factors. Comparisons with other ecotone movements offer further insights into the stability properties of the forest-savanna ecosystem. The boreal spread is widely understood to be a so-called pulled front, where the forest is the only stable solution for a set of environmental conditions (Abis & Brovkin, 2017). In that context, we are seeing a range expansion due to warming climate (Rees *et al*., 2020), which allows trees to survive where it was previously not possible. It is in this case that we see a fast spread, of up to 100 [m/yr]. In contrast, the dynamics of dryland shrub encroachment appears to be quite slow (Drees *et al*., 2023), less than 1 [m/yr]. Shrub encroachment is one of the famous examples of an apparent bistabilty of the system (D’Odorico *et al*., 2012), leading to pushed fronts (contrasted with pulled fronts, as previously noted). Given that, growth and spread dynamics of the dominant plants in all these systems are relatively similar, one can hypothesize that these plant species are capable of faster spread rates (e.g. 100 [m/yr]), but are slowed down due to bistable dynamics. If this is the case, then despite the relatively fast rates we see in our forest-savanna system (of the order of 5 [m/yr], Fig. 3), it is still more likely a bistable system. This gives further insight into the savanna question, of whether savanna-forests mosaics are may be bistable.

Parameter estimation by inference from dynamical data, is not widely found in the literature, and mainly restricted to species invasions and epidiomology (Keeling *et al*., 2004; Soubeyrand & Roques, 2014). This is likely due to lack of high-resolution spatio-temporal data, but this reality is changing with an explosion of widely available high-resolution remote sensing tools. To our knowledge, this study is the first instance of such an inference in the context of ecotone movement and encroachment dynamics. Overall the method works well to give a general estimate, but without more patterns (e.g. not only spread speed) or better estimates of other parameters, it is not very accurate.

One often overlooked topic in the context of estimation and modeling of such spread dynamics is the human dimension. Indeed, incorporating human impacts within ecological frameworks remains a challenge for modeling efforts in these complex landscapes. In the context of forest-savanna mosaics, several factors are likely to be most relevant. The spatial distribution and the temporal regime of ignition is strongly related to human activities. They also less directly modulate wild fires spread, for instance through grass fuel removal by grazing, which has also been evidenced at large scale by wild grazers in Serengeti (McNaughton, 1992). Both ignition and spread serves as critical factor in understanding landscape dynamics and ecosystem resilience. Conversely, small-scale logging in the forest patches may create opportunities for wild fire occurrences deep inside the forest patches. Integrating such human impacts into ecological models represents a crucial step towards developing comprehensive conservation and management strategies.

In conclusion, the findings presented in this study provide valuable insights into the complex dynamics of forest encroachment and spread movements in terrestrial ecosystems. By advancing our understanding of ecological processes, this research contributes to the development of more informed conservation and management strategies tailored to dynamic landscapes.

## A Appendix: Notes on inference

### A.1 Inference of grass diffusion coefficient

As noted in the main text, we could not directly estimate the diffusion parameter of grass (*D*_*G*_) from our empirical observations. This is because we did not observe cases of forest retreat, and with only forest encroachment cases, the ability to infer *D*_*G*_ is greatly diminished – the effect of *D*_*G*_ on front speed when forest is advancing is typically of the order of 0.1 [m/sec], which is much smaller than the actual variability of front speed (of the order of 10 [m/sec]). This means that front speed estimates simply do not contain enough information for inference of *D*_*G*_.

As a rough substitute, we look at historical data on the region, as analyzed by Lebamba *et al*. (2016). We use their observations, combined with extrapolation of savanna and forest spatial extent in Mpem and Djim National Park (MDNP), to arrive at two possible scenarios of historical savanna spread, which we term fast and slow.

In these scenarios the typical speed of grass spread rates are 0.25 [m/yr] and 2.65 [m/yr], for the slow and fast scenarios, respectively. Lebamba *et al*. (2016) find that the savanna has spread into the forest during the last few millennia. Based on their estimates (e.g. Fig.5 in Lebamba *et al*. (2016)), we can arrive at a slow spread scenario that took 3200 years, and a fast spread scenario of 1000 years. Inferring the possible state of MDNP over this period of time, we can use image morphological operations (Chudasama *et al*., 2015) to arrive at different fractions of savanna spread. We use a slow spread scenario of 50% savanna to 80% savanna, and a fast spread scenario of 5% savanna to 95% savanna, and in both cases ignore the 2% of the least reachable savanna pixels. From this we can estimate typical grass front spread rates of 0.25 [m/yr] and 2.65 [m/yr], for the slow and fast spread scenarios, respectively. We build front speed distributions with these values as median values, based on log-normal distributions that mimic the distributions shown in Fig. 3. For these log-normal distributions we use a *σ* value of 0.2 (i.e. the Gaussian distribution with *σ* = 0.2, and then apply a logarithm on it).

### A.2 Dependence on model stability

In the main text, for simplicity and generality, we do not consider the stability properties of the non-spatial states of the model when making the inference. As noted, given the 1000 parameter sets we randomly choose, this gives us 869 parameters sets where both forest and savanna states exists (and in practice, where at least one of them is stable). We compare here the inference results (i.e. in left panel of Fig. 4) for this set to that of a more constrained group of parameter sets, where both solutions are stable. In this case we have 186 parameter sets, and as seen in Fig. S1, results in a similar inference (right panel).

**Figure S1:**
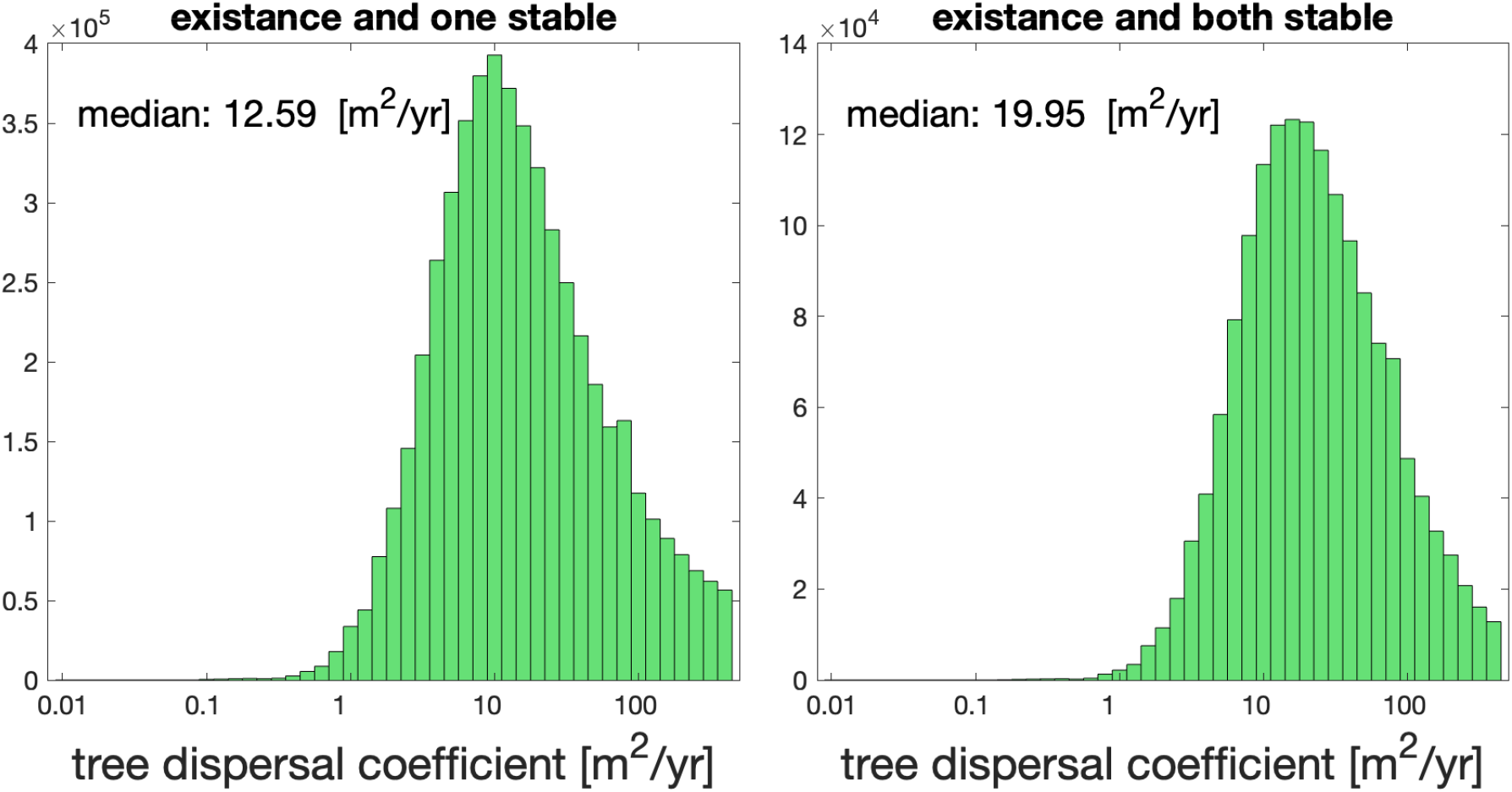
Inference of model parameters: diffusion coefficient *D*_*T*_. Left panel corresponds to the left panel of Fig. 4 in the main text, whereas right panel does the same analysis for a more constrained group of parameter sets, of only those where both forest and savanna solutions are stable.

